# Investigating visuo-tactile mirror properties in Borderline Personality Disorder: a TMS-EEG study

**DOI:** 10.1101/2024.05.29.596200

**Authors:** Zazio Agnese, Lanza Cora Miranda, Stango Antonietta, Guidali Giacomo, Marcantoni Eleonora, Lucarelli Delia, Meloni Serena, Bolognini Nadia, Rossi Roberta, Bortoletto Marta

**Affiliations:** Neurophysiology Lab, IRCCS Istituto Centro San Giovanni di Dio Fatebenefratelli, Brescia (Italy); Department of Psychology and Milan Center for Neuroscience - NeuroMI, University of Milano-Bicocca, Milan (Italy); Center for Cognitive Neuroimaging, School of Neuroscience and Psychology, University of Glasgow (UK); Psychiatry Unit, IRCCS Istituto Centro San Giovanni di Dio Fatebenefratelli, Brescia (Italy); Laboratory of Neuropsychology, IRCCS Istituto Auxologico Italiano, Milan (Italy)

**Author notes:** Center for Mind/Brain Sciences-CIMeC, University of Trento, Rovereto, (Italy) (Lanza Cora Miranda). Department of Neuroscience, Imaging and Clinical Sciences, University of Chieti-Pescara, Chieti (Italy) (Lucarelli Delia).

**Keywords:** empathy, cross-modal integration, TMS-evoked potentials, tactile mirror system, connectivity, psychiatric disorders, preregistered

## Abstract

Patients with Borderline Personality Disorder (pw-BPD) are characterized by lower levels of cognitive empathy compared to healthy controls (HCs), indicating difficulties in understanding others’ perspective. A candidate neural mechanism subtending empathic abilities is represented by the Tactile Mirror System (TaMS), which refers to mirror-like mechanisms in the somatosensory cortices. However, little is known about TaMS alterations in BPD, specifically in terms of brain connectivity within this network. Here, we aimed at providing novel insights on TaMS as neurophysiological candidate for BPD empathic deficits, with a special focus on TaMS connectivity by means of the combined use of transcranial magnetic stimulation and electroencephalography (TMS-EEG). Twenty pw-BPD and 20 HCs underwent a thorough investigation: we collected measures of empathic abilities obtained from self-report questionnaires, behavioral performance in a visuo-tactile spatial congruency task, and TMS-evoked potentials (TEPs) as effective connectivity indexes. In the TMS-EEG session, TMS was delivered over the right primary somatosensory cortex (S1) following the presentation of real touches and visual touches, while 74-channel EEG was continuously recorded. In the visuo-tactile spatial congruency task and the TMS-EEG recording, control conditions with visual touches on objects instead of body parts enabled to disentangle the involvement of TaMS from non-specific effects. The study is the first one employing TMS-EEG in pw-BPD and it has been preregistered before data collection. Consistent with previous findings, results show that pw-BPD reported significantly lower levels of cognitive empathy. Moreover, pw-BPD made significantly more errors than controls in the visuo-tactile spatial congruency task during visual touches on human body parts and not on objects. Finally, pw-BPD displayed a different connectivity pattern from S1-TEPs that was not specific for TaMS: they showed a lower P60 component during touch observation, as well as reduced amplitude of later TEPs responses (after ∼100 ms) during real touches. Overall, the present study shows behavioral evidence of TaMS impairment and a more general alteration in the connectivity pattern of the somatosensory network in pw-BPD.

## Introduction

The psychiatric condition known as Borderline personality disorder (BPD) is characterized by difficulties in emotional and behavioral regulation, issues with self-image and alterations in interpersonal relationships. Indeed, many of the symptoms can be attributed to deficits in social cognition, including difficulties in mentalization and empathizing (Lazarus, Cheavens, Festa, & Zachary Rosenthal, 2014; Sosic-Vasic et al., 2019). Specifically, patients with BPD (pw-BPD) exhibit reduced levels of cognitive empathy in comparison to individuals without the disorder. This suggests challenges in understanding others’ perspective, while the affective dimension shows unaltered or even increased levels (i.e., sensing others’ feelings) (Grzegorzewski, Kulesza, Pluta, Iqbal, & Kucharska, 2019; Harari, Shamay-Tsoory, Ravid, & Levkovitz, 2010; Martin, Flasbeck, Brown, & Brüne, 2017). Given the detrimental effects empathic abilities have on social connections, it is imperative to comprehend the cognitive and neurophysiological underpinnings of this impairment.

Empathic responses have been linked to the activity of mirror neuron systems. In this view, the understanding of others’ actions and sensations, as well as intentions and emotions, takes place through automatic simulation processes (Keysers & Gazzola, 2009). From the original discovery of mirror neurons in the monkey’s premotor ventral area (di Pellegrino, Fadiga, Fogassi, Gallese, & Rizzolatti, 1992; Gallese, Fadiga, Fogassi, & Rizzolatti, 1996), analogous mirror-like mechanisms in humans have been described both in the motor (Barchiesi & Cattaneo, 2015; Buccino et al., 2004; Catmur, Walsh, & Heyes, 2007; Ubaldi, Barchiesi, & Cattaneo, 2015) and in the somatosensory domain (Blakemore, Bristow, Bird, Frith, & Ward, 2005; Schaefer, Xu, Flor, & Cohen, 2009). In the so-called tactile-mirror system (TaMS), the same cortical areas involved in tactile perception are activated during the observation of others being touched (Keysers, Kaas, & Gazzola, 2010; Pihko, Nangini, Jousmaki, & Hari, 2010). Importantly, several neuromodulatory and neuroimaging studies highlighted the primary somatosensory cortex (S1) as a key area of the TaMS (Bolognini, Rossetti, Maravita, & Miniussi, 2011; Gazzola, Spezio, Etzel, Castelli, & Adolphs, 2012; Maddaluno, Guidali, Zazio, Miniussi, & Bolognini, 2020; Meyer, Kaplan, Essex, Damasio, & Damasio, 2011; Schaefer, Rotte, Heinze, & Denke, 2013; Zazio, Guidali, Maddaluno, Miniussi, & Bolognini, 2019). Interestingly, such embodied simulation processes in the somatosensory domain have been associated with empathy for pain (Lamm, Decety, & Singer, 2011) and with levels of cognitive empathy in healthy subjects (Bolognini, Miniussi, Gallo, & Vallar, 2013; Bolognini, Rossetti, Fusaro, Vallar, & Miniussi, 2014). Moreover, several studies reported that BPD patients showed abnormal processing of somatosensory stimuli, mainly in terms of nociception (Bohus et al., 2000; Schmahl & Baumgärtner, 2015; Schmahl et al., 2006), but also in tactile sensitivity as well as affective touch (Cruciani et al., 2023).

The neurophysiological alterations that may subtend the impairment in cognitive empathy typical of BPD are far from being understood. So far, neuroimaging studies have shown alterations in the mirror-like systems within the sensori-motor areas in BPD; however, findings are mixed, reporting either hyper-(Sosic-Vasic et al., 2019) or hypo-activations (Mier et al., 2013). Crucially, possible alterations of functional connectivity within these systems have not been investigated yet. While previous research described abnormal microstructural and functional brain connectivity in BPD (Orth, Zweerings, Ibrahim, Neuner, & Sarkheil, 2020; Quattrini, Marizzoni, et al., 2019; Quattrini, Pini, et al., 2019; Shafiei et al., 2024), evidence of connectivity alterations within the mirror systems in general, or in the TaMS in particular, is lacking. In this context, the combined use of transcranial magnetic stimulation and electroencephalography (TMS-EEG) has proven to be a promising tool to understand network dynamics, specifically in terms of effective connectivity: the TMS allows to directly activate a cortical area, while EEG traces the spread of cortical activation from the stimulated area to connected ones (Miniussi & Thut, 2010; Momi et al., 2021; Zazio, Miniussi, & Bortoletto, 2021). Interestingly, TMS-EEG has been recently employed to investigate the TaMS in healthy subjects (Pisoni, Romero Lauro, Vergallito, Maddaluno, & Bolognini, 2018), but to the best of our knowledge it has never been applied in pw-BPD.

Here we aim at investigating empathic abilities and TaMS functioning in BPD, both in terms of behavioral performance and neurophysiological measures of brain connectivity. In a group of pw-BPD and healthy controls (HCs), we collected measures of: (i) empathic abilities, evaluated by means of self-report questionnaires, (ii) performance in a behavioral task involving TaMS activity, namely the visuo-tactile spatial congruency (VTSC) task adapted from Bolognini et al. (2014), and (iii) connectivity indexes, obtained by TMS-EEG recording and represented by TMS-evoked potentials (TEPs). The study has been preregistered on Open Science Framework (OSF) before data collection to improve scientific reproducibility (OSF link). Preregistered hypotheses were the following:

*(i) Reduced empathic abilities in the cognitive domain*. Based on previous findings (Grzegorzewski et al., 2019; Harari et al., 2010; Martin et al., 2017), we expected lower cognitive empathy in pw-BPD compared to HCs. Considering that the literature is less consistent regarding affective empathy, we made no a-priori hypothesis on the comparison between groups in this dimension.
*(ii) Reduced behavioral interference for TaMS activation*. The VTSC task presented the participants with a visual touch on either one of two hands depicted on the screen, and a real touch on one of the participants’ hands, with the two stimuli being spatially congruent or incongruent. Participants were asked to report as fast as possible on which hand they felt the real touch. The rationale of this task was that seeing body parts being touched should activate S1 through the TaMS and therefore negatively impact performance in incongruent trials in terms of RTs. In pw-BPD, consistently with the hypothesis of a lower cognitive empathy, we expected a reduced interference effect compared to HCs, represented by a reduced impact of spatial incongruency of visual-touch when directed on human body parts. Moreover, in HCs we expected the interference effect in incongruent trials to be greater during visual touches on body parts than in a control condition with visual touches on objects. Finally, in HCs we hypothesized a negative relationship between cognitive empathy and performance in the VTSC in terms of RT, such that the greater the empathic abilities, the greater the interference effect (Bolognini et al., 2013, 2014).
*(iii) Altered connectivity pattern during TaMS activation*. Based on previous findings (Bolognini et al., 2014; Maddaluno et al., 2020), we hypothesized that 150 ms could represent the time interval needed for S1 to be activated through cross-modal integration within the TaMS network. Therefore, in the TMS-EEG session, TMS pulses were delivered 150 ms after a visual touch, and we expected a difference between pw-BPD and HCs when the visual touch was directed on human body parts only. We had no a-priori hypothesis on the direction of this effect, since both hyper-(Sosic-Vasic et al., 2019) and hypo-activations (Mier et al., 2013) of TaMS have been reported in the literature on BPD, although with different methodologies.

Moreover, to rule out the possibility that pw-BPD and HCs show differences in the somatosensory reafference (which does not involve the TaMS), the two groups were compared in TEPs obtained when a real touch stimulus was delivered in association with a TMS pulse with a time interval of 20 ms between the two stimuli, which should represent the time interval needed for S1 to be activated from peripheral reafference (Cohen, Bandinehi, Sato, Kufta, & Hallett, 1991).

## Materials and Methods

### Sample size estimation

Up to date, the literature does not provide studies in pw-BPD using performance at the VTSC task and TMS-EEG measures as dependent variables. Therefore, for these measures the present work represents a pilot study. For the sample size estimation we focused on the comparison between pw-BPD and HCs in the empathic levels, considering the works by Harari et al. 2010 and Martin et al. 2017. Sample size has been estimated using G*Power (3.1.9.7), considering a power of 80% and a threshold for statistical significance of 0.05. The results by Harari and colleagues (2010) indicate a significant Group (pw-BPD and HCs) X Empathy (Cognitive Empathy, Affective Empathy) interaction (F(1,40) = 6.38, *p =* 0.016), with pw-BPD showing a significantly lower cognitive empathy compared to HCs, resulting in a sample size of 12 participants per group. In the work by Martin et al. 2017, pw-BPD show lower cognitive empathy compared to HCs (*t*(41) = -3.78, *p* < 0.01), resulting in a sample of 20 participants per group. Taken together, we considered the estimation of the biggest sample size, i.e. 20 pw-BPD and 20 HCs.

### Participants

Twenty-three pw-BPD and 21 HCs were enrolled in the study after giving written informed consent. All participants were right-handed according to the Edinburgh Handedness Inventory (Oldfield, 1971), and with no contraindication to TMS (Rossi et al., 2021); they received a monetary reimbursement for travel expenses. Overall, 4 participants (3 pw-BPD, 1 HCs) did not take part in the TMS-EEG session as the rMT exceeded 82% of the maximal stimulator output (MSO; see exclusion criteria). The final sample consisted of 20 pw-BPD (3 men, mean age ± SE: 22.1 ± 0.8 years, range: 18-30 years) and 20 HCs (3 men, mean age ± SE: 23.3 ± 0.9 years, range 20-33 years). 18/20 pw-BPD were under pharmacological treatment.

The study was performed in accordance with the ethical standards of the Declaration of Helsinki and approved by the Ethical Committee of the IRCCS Istituto Centro San Giovanni di Dio Fatebenefratelli (Brescia, 65/2020).

### Clinical assessment

Adult patients meeting the criteria for inclusion in the study were selected based on a clinical diagnosis of BPD in accordance with the DSM-5 guidelines. The screening process was carried out by the Research Unit of Psychiatry of the IRCCS Fatebenefratelli, using comprehensive psychological evaluation that incorporated the use of the Structured Clinical Interview for DSM-5 Personality Disorders (SCID-5) to ascertain the presence of BPD. Patients were not included in case of comorbidity with schizophrenia and other psychotic disorders, according to DSM-5, and in case of unstable pharmacological therapy.

The severity of the BPD symptoms were assessed using the Zanarini rating scale for BPD (ZAN-BPD, (Zanarini, 2003) and the general state-psychopathology with the Symptoms Check-list 90 Revised (SCL-90-R, (Derogatis, 1994)). Depressive symptoms were evaluated with the Beck Depression Inventory II (BDI-II, (Beck, 1988)), impulsiveness with the Barratt Impulsiveness Scale (BIS, (Patton, Stanford, & Barratt, 1995)), and alexithymia with the Toronto Alexithymia Scale (TAS-20, (Bagby, Taylor, & Parker, 1994)). Moreover, interpersonal functioning was evaluated with the Interpersonal Problems (IIP, (Pilkonis, Kim, Proietti, & Barkham, 1996)), and attachment style will be assessed with the Attachment Style Questionnaire (Feeney, Noller, & Hanrahan, 1994). Finally, the Childhood Trauma Questionnaire (CTQ; (Bernstein & Fink, 1998)) was administered for the assessment of traumatic experiences, and the Inventory of statements about self-injury (ISAS, (Klonsky & Glenn, 2009)) for the evaluation of self-harm.

### Design and procedures

Participants underwent two experimental sessions on separate days. During Session 1, they performed the VTSC task, while in Session 2 they underwent the TMS-EEG recording. The two self-assessment questionnaires on empathic levels were completed at the end of Session 2, both for HCs and pw-BPD.

### Self-report questionnaires

Empathic response was measured by means of two self-report questionnaires, assessing cognitive and affective empathy: the Questionnaire of Cognitive and Affective Empathy (QCAE) ((Reniers, Corcoran, Drake, Shryane, & Völlm, 2011); Italian version: (Di Girolamo, Giromini, Winters, Serie, & de Ruiter, 2019)) and the Interpersonal Reactivity Index (IRI) ((Davis, 1983); Italian version: (Albiero, Ingoglia, & Alida, 2006)), both already administered in pw-BPD in previous works (Grzegorzewski et al., 2019; Harari et al., 2010; Martin et al., 2017). The QCAE consists of 31 statements, divided into 5 subscales (Perspective Taking, Online Simulation, Emotion Contagion, Proximal Responsivity, Peripheral Responsivity), rated on a 4-point Likert scale from 1 (“strongly agree”) to 4 (“strongly disagree”). The IRI consists of four 7-item subscales (Perspective Taking, Fantasy, Empathic Concern, Personal Distress) rated on a 5-point Likert scale from 0 (‘‘that does not describe me well”) to 4 (‘‘that describes me very well”).

### Visuo-tactile spatial congruency (VTSC) task

The VTSC task was adapted from Bolognini et al. (2014), which, in turn, referred to (Banissy & Ward, 2007). Participants were comfortably seated at 75 cm from a computer monitor (LCD, resolution 1280×800, refresh rate 60 Hz; head position ensured using a chinrest), on which they were presented with a left and a right hand in an egocentric perspective (stimuli eccentricity: 8° visual angle - Hands block). In each trial (**Figure 1A**), another hand in an allocentric perspective appeared on the top of the screen, and moved towards either the left or the right hand by means of four 100 ms-frames; the final frame (1000 ms duration) shows the allocentric hand touching the one of the egocentric hands (visual touch). After 10 ms from the beginning of the visual touch, a real tactile stimulus (real touch) was delivered either on participants’ left or right hand, administered through miniature solenoid tappers (visuo-tactile trials). Visual and real touches were spatially congruent or incongruent. In unimodal trials, only the visual touch (visual-only) or the real touch (tactile only) were presented. Participants were asked to fixate a red asterisk in the center of the screen and to report the side of the real touch by button press on a computer keyboard. Performance was evaluated in terms of accuracy and reaction times (RTs). In catch-no-touch trials, the color of the fixation cross changed to green, and no touches (neither visual nor real) were presented. In these trials, participants were asked to press both the response buttons. Catch-no-touch trials ensured that participants maintained the fixation at the center of the screen. In a control block, the two hands in the egocentric perspective were replaced with two objects (i.e., two leaves - Leaves block). Block order was counterbalanced across participants.

**Figure 1.**
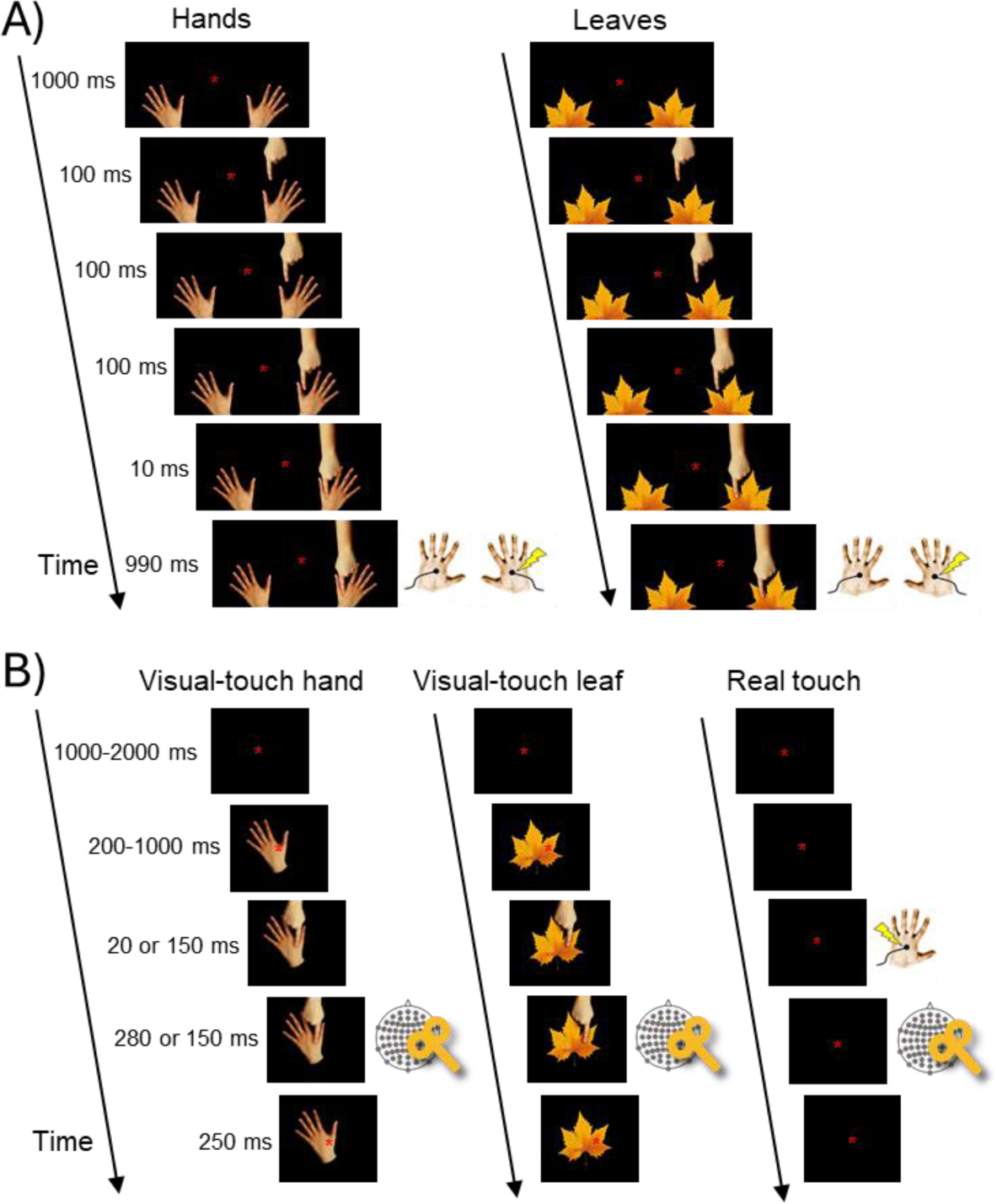
Schematic representation of trials of the VTSC task and the TMS-EEG recording. **A)** VTSC task. Example of a congruent trial with visual touches on the right during the Hands block (left) and the Leaves block (right), showing the visual frames of the approaching hand towards the hands (or the leaves) with relative temporal durations, and the real touch delivered 10 ms after visual-touch onset. In incongruent trials with the same visual touches on the right, real touch was delivered on the left hand. Analogous trial types were presented also for visual touches on the left, and block order was randomized. **B)** TMS-EEG recording. Representation of TMS-trials during the presentation of touches (i.e., visual-touch on the hand, visual-touch on the leaf, real touch), with the TMS pulse over S1 delivered 20 or 150 ms after visual touch or real touch onset. In half of the trials, TMS was not delivered; trial order was randomized. In both A) and B), reported time intervals represent frame durations.

At the end of each block, participants were asked to report on a brief questionnaire their sensations by using a visual analog scale (VAS) in each item; the questionnaire comprised the following items: (1) “When I was shown with a *hand/leaf* being touched, I had the feeling of being touched on my own hand (2) “Looking at the *hand/leaf* being touched made it difficult to localize the tactile stimulus on my own hand”.

The experiment was run in E-Prime software (E-Prime 2.0, Psychology Software Tool, Inc.). Timing of stimuli delivery has been checked by using a photodiode (for visual stimuli) and a pressure sensor (for tactile stimuli).

### TMS-EEG

EEG signal was recorded at a sampling rate of 9600 Hz from 74 TMS-compatible passive Ag/AgCl electrodes (EasyCap, BrainProducts GmbH, Munich, Germany) by means of a TMS-compatible system (g.HIamp, g.tec medical engineering GmbH, Schiedlberg, Austria). No filters were applied during the recording and the skin-electrode impedance was kept below 5 kΩ.

TMS pulses were delivered using a Magstim Rapid2 stimulator (Magstim Company, Whitland, UK) with a 70 mm figure-of-eight coil (Alpha B.I.), which produced a biphasic waveform. The orientation of the coil was approximately 45° from the midline so that the direction of current flow in the right S1 during the second phase of the pulse waveform was posterior-to-anterior (Siebner et al., 2022; Sommer et al., 2006). The charge delay was set at 350 ms and coil position was monitored using the Softaxic 3.4.0 neuronavigation system (EMS, Bologna, Italy). To attenuate the contamination of TEPs with sensory artifacts, participants wore noise-canceling earphones playing white noise and a thin layer of foam was applied under the coil.

First, the motor hotspot for the left First Dorsal Interosseous (FDI) muscle was localized as the scalp location eliciting the highest and most reliable motor-evoked potentials (MEPs) with the same TMS intensity. Then, the rMT was estimated using the maximum-likelihood threshold hunting algorithm (Awiszus, 2003, 2011), a variant of the best parameter estimation by sequential testing (best PEST) procedure (Pentland, 1980). MEPs during rMT estimation were not recorded. Once the individual rMT was determined, the location of the right S1 was identified by moving the TMS coil 2 cm lateral and 0.5 cm posterior to the hotspot for FDI (Holmes & Tamè, 2019). TMS intensity was set at 110% of rMT.

During the TMS-EEG session, participants were presented with real touches, delivered on the left hand by means of a solenoid tapper, and with visual touches, presented in the center of a computer screen (the same as the VTSC task, placed at 75 cm from participants; head position was ensured by using a chinrest). Visual touches consisted of a left hand or a leaf being touched. Both TMS trials and no-TMS trials were included. In TMS trials, a TMS pulse was delivered over participants’ right S1 after the visual or the real touch stimulus. The inter-stimulus interval (ISI) between the visual/real touch stimulus and the TMS was either 20 ms or 150 ms (**Figure 1B)**. No-TMS trials were identical to TMS trials except that TMS was not delivered after the visual/real touch stimuli. Based on pilot data (see Preregistration on <u>OSF</u>), these trials were used to extract the event-related potentials (ERPs) to the visual and real touch stimuli. For each of the 9 conditions (TMS trials: 3 x 2, Trial type X ISI; No-TMS trials: 3 Trial type), 77 trials were presented, divided in 7 blocks; trial order was randomized. To maintain participants’ attention on the visual stimuli, 28 catch trials requiring participants’ response were included. TMS was not delivered during catch trials. The number of trials has been determined to obtain a good signal-to-noise ratio in TEPs, based on previous TMS-EEG studies (e.g., Bortoletto et al. 2021) and on the pilot experiment, while keeping the total duration of the experiment as short as possible, considering the involvement of a clinical population.

The experiment was run in E-Prime software (E-Prime 2.0, Psychology Software Tool, Inc.). Timing of stimuli delivery has been checked by using a photodiode (for visual stimuli) and a pressure sensor (for tactile stimuli).

### Exclusion criteria

Participants were excluded from the sample in the following cases: (i) they did not complete all blocks of the VTSC session; (ii) performance at catch trials (‘catch-no-touch’ trials in the VTSC and ‘catch’ trials in the TMS-EEG session) was below 50%; (iii) in the VTSC session, RTs or number of errors in at least one condition deviated of more than 2.5 SD from the sample mean; (iv) in the TMS-EEG session, 110% of rMT exceeded 90% of the maximal stimulator output; (v) in the TMS-EEG session, the final TEPs obtained for each trial type comprised less than 54 trials (i.e., 70% of the planned 77 trials) (**Figure 2A-B**).

**Figure 2.**
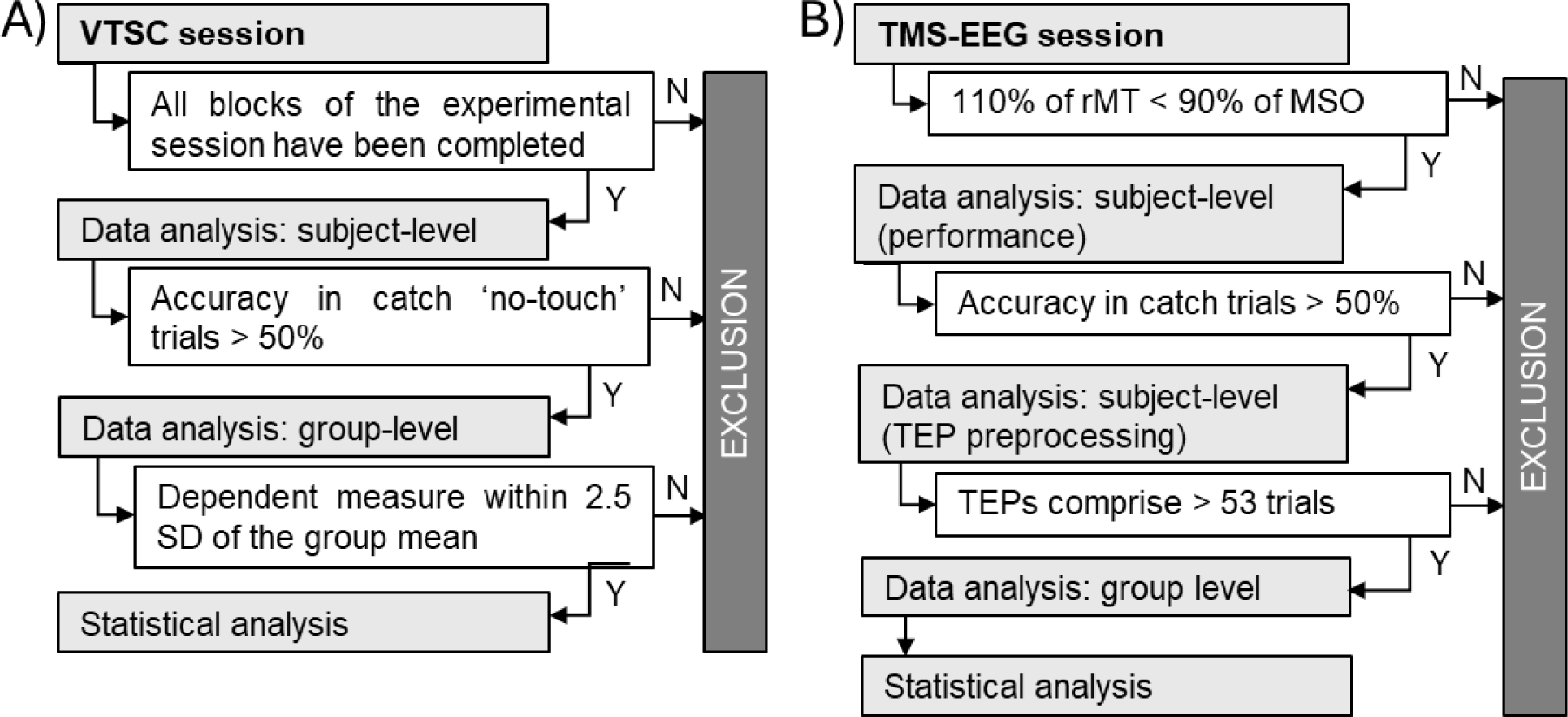
Preregistered exclusion criteria for VTSC session (A) and TMS-EEG session (B).

## Analysis

### Self-report questionnaires

For the QCAE, the measure for Cognitive Empathy was obtained by the sum of the scores in the Perspective Taking and Online Simulation subscales, while for Affective Empathy by the sum of the scores in the Emotion Contagion, Proximal Responsivity and Peripheral Responsivity. For the IRI, the measure for Cognitive Empathy was obtained by the sum of the scores in the Perspective Taking and Fantasy subscale, while for Affective Empathy by the sum of the scores in the Empathic Concern and Personal Distress subscales. Normative data for the Italian population (Maddaluno, Aiello, Roncoroni, Prunas, & Bolognini, 2022) were applied to raw values for exploratory analyses.

### Behavioral data

Individual RTs data was log10-transformed, and trials exceeding ± 2 SD of the individual mean were discarded. Then, the average value for each trial type was considered for the statistical analyses. Accuracy was measured in terms of number of errors.

### TMS-EEG data

One pw-BPD was excluded from the analyses as she did not complete all TMS-EEG blocks due to discomfort during TMS delivery, and the minimum number of trials was not reached (see exclusion criteria). Thus, a sample of 20 HCs and 19 pw-BPD was left for ERP and TEP analysis. TMS-EEG data processing was performed in MATLAB R2020b (The Mathworks, Natick, MA, USA) with custom scripts using EEGLAB v.2020.0 (Delorme & Makeig, 2004) and FieldTrip functions (Oostenveld, Fries, Maris, & Schoffelen, 2011), and following the same steps applied in previous studies by our research group (Guidali, Zazio, et al., 2023; Zazio, Barchiesi, Ferrari, Marcantoni, & Bortoletto, 2022). Unless otherwise specified, the default parameters for the EEGLAB and FieldTrip functions were used.

For each participant, the first pre-processing steps included all trial types. This procedure ensured that the same pre-processing steps were applied to all conditions, thus minimizing the risk of differences between conditions arising from dissimilarities in the preprocessing steps. Continuous TMS-EEG data was interpolated for 3 ms around the trigger to eliminate TMS pulse induced artifact, high-pass filtered at 1 Hz (FIR sync filter, EEGLAB function ’pop_eegfiltnew’, order 31682), downsampled to 4800 Hz and epoched from -750 ms before to 750 ms after the stimulus. Subsequently, the source-estimate-utilizing noise-discarding (SOUND) algorithm was applied to discard noise measurement (spherical 3-layer model, regularization parameter: λ=.01; Mutanen et al., 2018), followed by a first round of automatic artifact rejection (EEGLAB function ‘pop_jointprob’, threshold for rejection: 5 SD). Then, ocular artifact correction was performed by means of Independent Component Analysis (ICA; EEGLAB function ’pop_runica’, infomax algorithm, 73 channels included, 73 ICA components calculated): the horizontal and vertical eye movement components were visually inspected. Then, the signal-space projection and source-informed reconstruction (SSP-SIR, Mutanen et al., 2016) algorithm was applied for the removal of the TMS-evoked muscle artifact in the first 50 ms after the TMS pulse; principal components were visually inspected and discarded if they represented a high-frequency signal time-synchronized with the TMS pulse. Finally, the data was filtered with a 70 Hz low-pass filter (IIR Butterworth filter, order 4, EEGLAB function ’pop_basicfilter’) and re-referenced to average reference. Both ICA and SSP-SIR steps were performed by two independent researchers. Then, epochs were redefined around the trigger at a range of -200 ms to 400 ms and the baseline corrected to -200 ms to -2 ms, and a second manual artifact rejection was performed to discard residual artifactual trials. At this point, the data were divided according to trial type.

The same pipeline, except for the signal interpolation and SSP-SIR, was run for non-TMS trials to obtain the ERPs generated by the presentation of the visual stimuli (baseline) as well as ERPs generated by the visual-touch stimuli (i.e., real touches and visual touches on the hand or on the leaf).

After preprocessing, the data was converted into FieldTrip structures for visualization and statistical analysis.

In the exploratory analysis on TEP peaks, we focused on the early components within 100 ms to avoid confounds related to the TMS sensory processing (Herring, Esterer, Marshall, Jensen, & Bergmann, 2019; Niessen, Bracco, Mutanen, & Robertson, 2021; Nikouline, Ruohonen, & Ilmoniemi, 1999). Peaks’ amplitude and latency were extracted from electrode CP4, which showed the highest signal in all components in the mean of all conditions, by averaging over 10 ms around the peak.

### Statistical analysis

If not otherwise specified, the statistical analyses followed what was planned in the preregistration.

I. *Self-report questionnaires.* For each questionnaire (i.e., QCAE and IRI), we ran a 2 X 2 repeated-measure analysis of variance (rm-ANOVA) with within factor Type (Cognitive empathy, Affective empathy) and between factor Group (HCs, pw-BPD). Exploratory descriptive analyses included the application of IRI score correction based on Italian normative data (Maddaluno et al., 2022), to obtain equivalent scores for each participant.
II. *VTSC task.* As preliminary analyses, independent t-tests compared HCs and pw-BPD on accuracy at visual-only trials and on RT at tactile-only trials, to rule out the presence of generic group differences. In this way, possible differences between groups may be attributed to the activity of TaMS rather than to specific differences in reaction times. Then, on visuo-tactile trials, we run a 2 X 2 X 2 rm-ANOVA with within factors Stimulus (Hands, Leaves) and Congruency (Congruent, Incongruent), and between factor Group (HCs, pw-BPD). The rm-ANOVA was planned for both accuracy (i.e., number of errors) and RT. However, for accuracy, due to ceiling effects we decided to apply non-parametric tests: U Mann-Whitney instead of Student t-test for the preliminary analysis, and Friedman ANOVA on the Incongruent-Congruent difference (ΔACC(Incong-Cong)) instead of the 2 X 2 X 2 rm-ANOVA. Regarding the expected effects on HCs, first, a correlation tested the relationship between Cognitive Empathy scores and the interference effect at the VTSC, defined as the difference in RT between incongruent and congruent trials in the Hands block. Second, a one-tail t-test for dependent samples compared the interference effect in the Hands and the Leaves blocks. Exploratory analyses on RT included the comparison between visuo-tactile congruent, visuo-tactile incongruent and unimodal tactile-only trials by means of a 3 X 2 rm-ANOVA with within factor Trial type (congruent, incongruent, tactile-only) and between factor Group, to reveal whether visual-touch shortened or lengthened the RT in unimodal tactile-only trials. This analysis was performed separately for the Hands block and the Leaves block. Finally, we subtracted visuo-tactile trials (i.e., congruent and incongruent trials) from tactile-only trials, to obtain a normalized RT score (ΔRTnorm) based on individual RT. The ΔRTnorm entered the 2 X 2 X 2 Stimulus X Congruency X Group rm-ANOVA described above. Exploratory analyses on the responses to the questionnaire on the sensations induced by the VTSC were analyzed by means of Friedman ANOVA, separately for each question.
III. *TMS-EEG.* The preregistration included two analyses on TEPs: First, considering TEPs obtained from Visual touch (touch-hand and touch-leaf) trials with ISI-150 in HCs and pw-BPD, and testing for an interaction effect of a 2 X 2 mixed between-within-subjects design. To this end, TEPs from touch-leaf trials were subtracted from TEPs from touch-hand trials and then compared between HCs and pw-BPD by means of a two-tailed non-parametric cluster-based permutation test for independent samples over all channels and time points from 4 to 350 ms after the TMS pulse (Maris & Oostenveld, 2007). Second, the same statistics were applied to compare HCs and pw-BPD on TEPs obtained from Real touch trials with ISI-20. Analyses on ERPs were exploratory. ERPs were calculated both for no-TMS trials, i.e., generated by the presentation of the hand or the leaf before the touch occurred (baselineERPs), as well as the ones generated by the presentation of the visual or the real touch (touchERPs). Exploratory analyses on TEPs with ISI-20 and ERPs from visual-touch trials followed the same approach, where applicable: we tested the main effects of Stimulus and Group, and the Stimulus X Group interaction by means of two-tailed non parametric cluster-based permutation tests. Specifically, the main effect of Stimulus was tested by concatenating data from HCs and pw-BPD and then performing a t-test for dependent samples, while the main effect of Group was tested by averaging signals from Hand trials and Leaf trials, and performing a t-test for independent samples. As in the preregistered analysis on TEPs with ISI-150, the interaction was tested by subtracting signals in Leaf trials from signals in Hand trials. Cluster-based analyses on TEPs were performed over from 4 to 350 ms after the TMS pulse, while cluster-based analyses on ERPs started from 1 ms after stimulus onset; all analyses were performed over all channels. Finally, an exploratory analysis on the TEP-ERP difference was performed on the amplitude and latencies of peaks of the main components, which entered a 2 X 2 X 2 Stimulus X ISI X Group rm-ANOVA. In this case, the threshold for significance was corrected for the two peaks (i.e., 0.05/2=0.025). HCs and pw-BPD were also compared in the rMT by means of an independent t-test.

The threshold for statistical significance was set at *p*<0.05. The statistical analysis of the data obtained from the questionnaires and the VTSC task were performed in Jamovi (The jamovi project 2.3.21, 2021; R Core Team, 2020), while statistics on neurophysiological measures (i.e., TEPs and ERPs) was performed in MATLAB R2020b (The Mathworks, Natick, MA, USA) using FieldTrip functions (Oostenveld et al., 2011). Reported *p* values are corrected for multiple comparisons, where applicable (Tuckey correction for data from questionnaires, the VTSC task and TEP peaks, cluster correction for the analysis on TEPs and ERPs over all channels and time-points). For the cluster-based analysis, the reported time intervals for significant clusters are to be intended as approximate latencies (Sassenhagen & Draschkow, 2019).

## Results

If not reported otherwise, data were normally distributed according to the Shapiro-Wilk test, and mean ± SE are reported in parentheses.

### Clinical evaluation

### Self-report questionnaires

Overall, data from questionnaires indicated an impairment in cognitive empathy in pw-BPD. The results on the QCAE highlighted a significant Empathy x Group interaction (F(1,38)=12.9, *p*=0.001, *η^2^_p_*=0.25), with post-hoc analyses revealing lower Cognitive Empathy in pw-BPD compared to HCs (*t*=2.94, *p*=0.028; **Table 1**). We also observed a main effect of Empathy (F(1,38)=316.34, *p*<0.001, *η^2^_p_*=0.89), showing higher values for Cognitive Empathy (57.7 ± 1.6) compared to Affective Empathy (36.9 ± 0.85), and a significant main effect of Group (F(1,38)=4.20, p=0.047, *η^2^_p_*=0.10), with HCs (49.5 ± 1.48) showing overall greater empathic levels compared to pw-BPD (45.2 ± 1.48). For the IRI, the Empathy X Group interaction showed a trend towards statistical significance (F(1,38)=3.76, *p*=0.06, *η^2^_p_*=0.09), with pw-BPD showing lower scores in Cognitive Empathy compared to HCs (**Table1**). As for the QCAE, we observed a significant main effect of Empathy (F(1,38)=4.341, *p*=0.0.044, *η^2^_p_*=0.1), and post-hoc comparisons revealed higher levels of Cognitive Empathy (35.97 ± 1.44) compared to Affective Empathy (33.1 ± 1.17). The main effect of Group was not significant (F(1,38)=1.76, *p*=0.192, *η^2^_p_*=0.04). Based on the IRI normative values (Maddaluno et al., 2022), 7/20 pw-BPD were found to be below the threshold for normal range (i.e., equivalent score of 0), index of a defective score, and 3/20 pw-BPD showed a borderline score for normal range (i.e., equivalent score of 1) in at least one of the subscales. Among HCs, only 1/20 reported an equivalent score of 1.

**Table 1.** Test scores at the clinical assessment of pw-BPD. N indicates the number of participants who completed the questionnaires.

| | N | Mean | ± | SE |
| --- | --- | --- | --- | --- |
| Zanarini rating scale for Borderline Personality Disorder (ZAN-BPD) | 13 | 10.54 | ± | 3.6 |
| Baratt Impulsiveness Scale (BIS-11) | 14 | 70.29 | ± | 11.07 |
| Beck Depression Inventory (BDI-II) | 14 | 29.21 | ± | 13.99 |
| Toronto Alexithymia Scale (TAS) | 14 | 60.79 | ± | 13.05 |
| Symptom Checklist-90-R (SCL90) | 13 | 168.92 | ± | 78.73 |
| Inventory of Interpersonal Problems (IIP) | 14 | 2.08 | ± | 0.65 |
| Childhood Trauma Questionnaire (CTQ) | 15 | 51.27 | ± | 13.06 |
| Attachment Style Questionnaire (ASQ) | 14 |  |  |  |
| *Confidence* | | 23.71 | ± | 6.52 |
| *Discomfort with Closeness* | | 42.5 | ± | 4.78 |
| *Relationships as secondary* | | 18.36 | ± | 9.68 |
| *Need for Approval* | | 29.86 | ± | 7.59 |
| *Preoccupation with Relationships* | | 36.93 | ± | 6.03 |
| Inventory of statements about self-injury (ISAS) | 19 |  |  |  |
| *Presence of self-injury (%)* |  | 84.21% |  |  |

**Figure 2.**
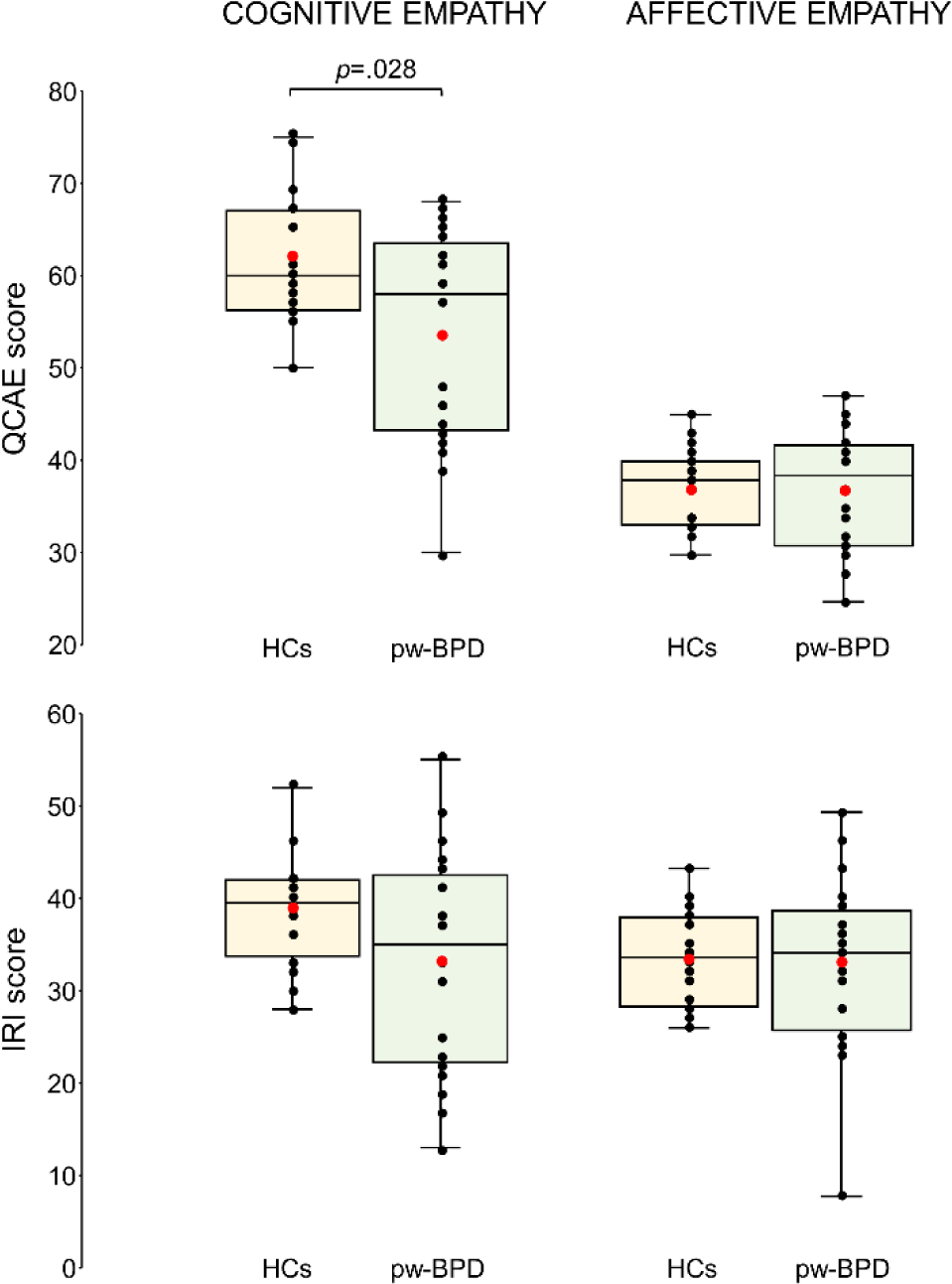
Results on the questionnaires for empathic abilities. *Upper panel:* QCAE; *lower panel*: IRI. In the box-and-whiskers plots, red dots indicate the means of the distributions. The center line denotes their median values. Black dots show individual participants’ scores. The box contains the 25^th^ to 75^th^ percentiles of the dataset. Whiskers extend to the largest observation, which falls within the 1.5 * inter-quartile range from the first/third quartile. The p-value of the significant results from Group X Empathy interaction is reported.

### VTSC

Taken together, results on the VTSC showed that pw-BPD performed worse than HCs in terms of accuracy (i.e, higher number of errors) when hand touches were presented, while RTs were affected by spatial congruency between the visual and the tactile stimuli in both groups.

Regarding accuracy, preliminary analyses showed that performance in Catch-no-touch trials was above 50% in all cases, indicating that participants were attending the visual stimuli; moreover, HCs and pw-BPD did not differ in the number of errors on visual-only trials (*U*=164, *p*=0.415), in which they were asked to not provide a response. Importantly, ΔACC(Incong-Cong) was significantly different across conditions in visuo-tactile trials (*χ2*=8.38, p=0.039), as indicated by a non-parametric Friedman ANOVA applied to account for deviation from normality (Shapiro-Wilk test p<0.001). Specifically, post-hoc pairwise Durbin-Conover comparisons revealed that pw-BPD made significantly more errors in the ΔACC(Incong-Cong) compared to HCs in the Hands block (*p*=0.016) but not in the Leaves block (*p*=0.36); remaining comparisons were not significant7.

In RTs, one HC exceeded ± 2.5 SD from the mean and therefore was excluded, leaving 19 HCs and 20 pw-BPD for VTSC analyses. The preliminary analysis showed no significant differences in RT in tactile-only trials between HCs and pw-BPD either in the Hand (*t*=-1.52, *p*=0.138) or in the Leaves block (*t*=-1.12, *p*=0.268), indicating that the two groups did not show non-specific differences on RTs. Importantly, the rm-ANOVA on visuo-tactile trials revealed a significant main effect of Congruency (*F(1,37)*=59.35, *p*<.001, *η^2^* =0.62), with slower RT in incongruent trials (mean ± SE: 2.56 ± 0.014) compared to congruent trials (mean ± SE: 2.54 ± 0.014). No other main effects (Stimulus: F=0.08, *p*=0.772, *η^2^*=0.002; Group: F=1.19, *p*=0.283, *η^2^* =0.03) nor interactions (Stimulus X Group: F=1.54, *p*=0.223, *η^2^* =0.04; Congruency X Stimulus: F=0.90, *p*=0.348, *η^2^* =0.02; Congruency X Stimulus X Group: F=0.006, *p*=0.940, *η^2^* =0.00) were significant. Mean values of raw RTs and error number are reported in **Table 2**.

**Table 2.** Descriptive data for pw-BPD and HCs. Demographics, average scores obtained at the different subscales of the QCAE and IRI questionnaires, average raw RTs and number of errors recorded during the different trial types of the VTSC task.

|  |  | HC |  |  | BPD |  |  |
| --- | --- | --- | --- | --- | --- | --- | --- |
|  |  | N(F)=20(17) |  |  | N(F)=20(17) |  |  |
|  |  | Mean | ± | SE | Mean | ± | SE |
| Age in years |  | 23.3 | ± | 0.9 | 22.1 | ± | 0.8 |
| Education in years |  | 14.6 | ± | 0.6 | 12.1 | ± | 0.7 |
| *QCAE* |  |  |  |  |  |  |  |
|  | Perspective Taking | 32.35 | ± | 0.96 | 28.79 | ± | 1.75 |
|  | Online Simulation | 29.5 | ± | 0.78 | 24.37 | ± | 1.71 |
|  | Emotion Contagion | 11.7 | ± | 0.56 | 12.42 | ± | 0.51 |
|  | Proximal Responsivity | 12.7 | ± | 0.39 | 12.05 | ± | 0.66 |
|  | Peripheral Responsivity | 12.55 | ± | 0.3 | 12.11 | ± | 0.74 |
|  | *Cognitive Empathy* | 61.95 | ± | 1.45 | 53.16 | ± | 2.66 |
|  | *Affective Empathy* | 36.95 | ± | 0.99 | 36.58 | ± | 1.46 |
| *IRI* |  |  |  |  |  |  |  |
|  | Perspective Taking | 19.3 | ± | 0.65 | 16.6 | ± | 1.59 |
|  | Fantasy | 19.5 | ± | 0.94 | 16.55 | ± | 1.57 |
|  | Empathic Concern | 20.7 | ± | 0.67 | 17.35 | ± | 1.3 |
|  | Personal Distress | 12.55 | ± | 0.76 | 15.6 | ± | 1.09 |
|  | *Cognitive Empathy* | 38.8 | ± | 1.26 | 33.15 | ± | 2.6 |
|  | *Affective Empathy* | 33.25 | ± | 1.12 | 32.95 | ± | 2.06 |
| *VTSC task - number of errors* |  |  |  |  |  |  |  |
|  | Visual touch congruent (Hands) | 0.15 | ± | 0.08 | 2.40 | ± | 1.42 |
|  | Visual touch incongruent (Hands) | 0.60 | ± | 0.15 | 3.95 | ± | 1.49 |
|  | Visual touch congruent (Leaves) | 0.25 | ± | 0.16 | 2.80 | ± | 1.22 |
|  | Visual touch incongruent (Leaves) | 0.65 | ± | 0.21 | 3.70 | ± | 1.13 |
|  | Tactile-only (Hands) | 0.90 | ± | 0.20 | 4.20 | ± | 1.81 |
|  | Tactile-only (Leaves) | 0.50 | ± | 0.33 | 4.15 | ± | 1.41 |
| *VTSC task - RTs* |  |  |  |  |  |  |  |
|  | Visual touch congruent (Hands) | 344.89 | ± | 13.82 | 384.97 | ± | 22.12 |
|  | Visual touch incongruent (Hands) | 355.68 | ± | 15.64 | 378.20 | ± | 22.97 |
|  | Visual touch congruent (Leaves) | 364.39 | ± | 13.20 | 415.13 | ± | 21.38 |
|  | Visual touch incongruent (Leaves) | 374.02 | ± | 15.90 | 404.19 | ± | 23.51 |
|  | Tactile-only (Hands) | 405.79 | ± | 15.94 | 452.13 | ± | 21.28 |
|  | Tactile-only (Leaves) | 412.75 | ± | 16.82 | 450.93 | ± | 23.26 |

Exploratory analyses did not reveal additional group differences, but an overall advantage in terms of RTs in visual-touch trials compared to unimodal tactile-only trials and a higher subjective sensation of being touched during the Hand block than during the Leaves block. Specifically, in the Trial type X Group rm-ANOVA on RTs, we observed a main effect of Trial type for both the Hands block (F(2,74)=84.74, p<0.001, *η^2^_p_* = 0.7) and the Leaves block (F(2,74)=43.94, p<0.001, *η^2^_p_*=0.54). Post-hoc comparisons showed longer RTs for unimodal tactile-only compared to both incongruent visuo-tactile trials and congruent visuo-tactile trials, and that in incongruent trials RTs were longer than congruent trials (p<0.001 in all comparisons). Main effect of Group was not significant either in the Hands block (F(1,37)=2.21, p=0.145, *η^2^_p_*=0.06) or the Leaves block (F(1,37)=0.99, p=0.327, *η^2^_p_*=0.03), as well as the Trial type X Group interaction (Hands: F(2,74)=0.42, p=0.658, *η^2^_p_*=0.01; Leaves F(2,74)=1.3, p=0.278, *η^2^_p_*=0.034). The analyses on the sensations induced by the VTSC highlighted a significant effect in the sensations described in Item-1 (*“When I was shown with a hand/leaf being touched, I had the feeling of being touched on my own hand”;* χ2=13.988 p=0.003). Post-hoc pairwise Durbin-Conover comparisons reveal that both pw-BPD and HCs reported higher scores for the Hands block compared to the Leaves block (HCs: *p*=0.031; pw-BPD: *p=*0.015). Item-2 (“Looking at the *hand/leaf* being touched made it difficult to localize the tactile stimulus on my own hand”) was not significant (χ2=1.817 p=0.611).

### Exploratory: Event-Related Potentials (ERPs)

In baselineERPs generated from the presentation of the hand or the leaf (i.e., before the visual touch occurred), ERPs features were affected by the Stimulus. We observed three significant clusters, two positive (*p*=0.03, from 70 ms to 110 ms over frontal and lateral right electrodes; *p*=0.002 from 100 ms to 200 ms over frontocentral electrodes), and one negative (*p=*0.002, from 70 ms on over posterior central electrodes). The N170, a component typically generated in response to images of faces and body parts (Kovács et al., 2006), was present in baselineERP(Hand) and not in baselineERP(Leaf) (**Figure 3**). No significant clusters were present, either for the main effect of Group (p=1) or for the Stimulus X Group interaction (p=1).

**Figure 3.**
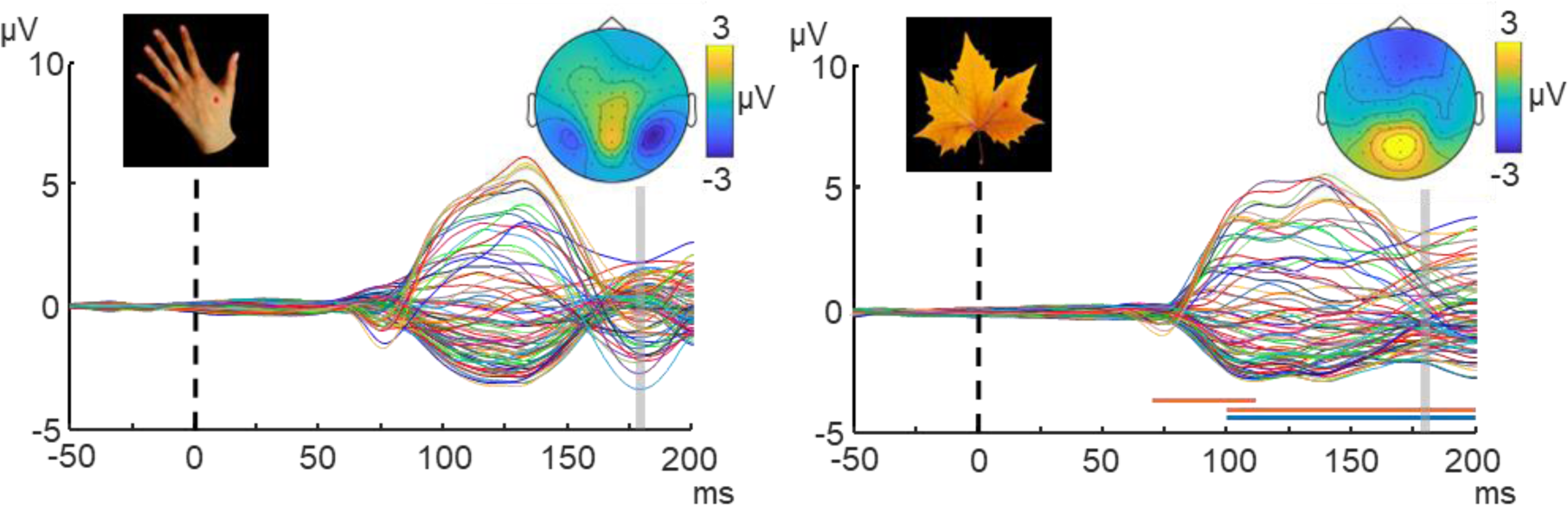
Main effect of Stimulus in baselineERPs. Butterfly-plot ERPs generated by the sight of a hand (left panel) and of a leaf (right panel). Topographies (amplitude range in colorbar) are obtained on the averaged signal between 160 and 180 ms after stimulus onset, and show the typical N170 pattern after the sight of a hand but not of a leaf. Horizontal lines indicate the latencies of significant clusters (orange: positive; blue: negative).

In touchERPs generated by the touch frame on the hand or on the leaf, results suggest a different pattern for pw-BPD and HCs depending on the visual stimulus. Indeed, the Stimulus X Group interaction showed a trend towards significance in one positive cluster (p=0.066). Exploratory direct comparisons between touchERP(Hands) and touchERP(Leaf) within each group reveal two significant clusters in pw-BPD, one positive (p=0.014) from 215 ms to 280 ms over frontal right electrodes and one negative (p=0.014) from 185 to 290 ms over posterior left electrodes, showing reduced ERPs component during human-directed touch compared to object directed touch. No significant clusters emerged in HCs (p>0.304). Finally, no significant clusters emerged for the main effect of Stimulus (p>0.102) or for the main effect of Group (p>0.436).

Real-touchERPs did not differ between HCs and pw-BPD, as no significant clusters emerged (p>0.753).

### TMS-Evoked Potentials (TEPs)

Preliminary analyses showed that during TMS-EEG accuracy in catch trials was always above 50%, indicating that all participants attended the visual stimuli; moreover, the rMT did not differ between HCs (mean ± SE: 61.1 ± 2.1) and pw-BPD (mean ± SE: 60.5 ± 2.5; p=0.856).

Preregistered analysis showed that TEPs were modulated by the content of the visual stimulus when TMS was delivered 150 ms after touch onset, but did not show differences between groups indicating specific TaMS alterations in pw-BPD. Indeed, in visual-touch trials with ISI-150 we observed two significant clusters for the main effect of Stimulus, one positive over posterior electrodes (p=0.002) and one negative over fronto-central electrodes (p=0.002), both from approximately 230 ms to 350 ms, resulting in reduced TEPs components during touches on the leaf compared to touches on the hand (Figure 4). No clusters emerged for the main effect of Group (p>0.142) nor the Stimulus X Group interaction (p>0.548). Instead, no significant effects were found in exploratory analyses on TEPs with ISI-20 on visual-touch trials, neither for the main effect of Stimulus (p>0.19), nor for the main effect of Group (p>0.18), nor for the Stimulus X Group interaction (p>0.59).

**Figure 4.**
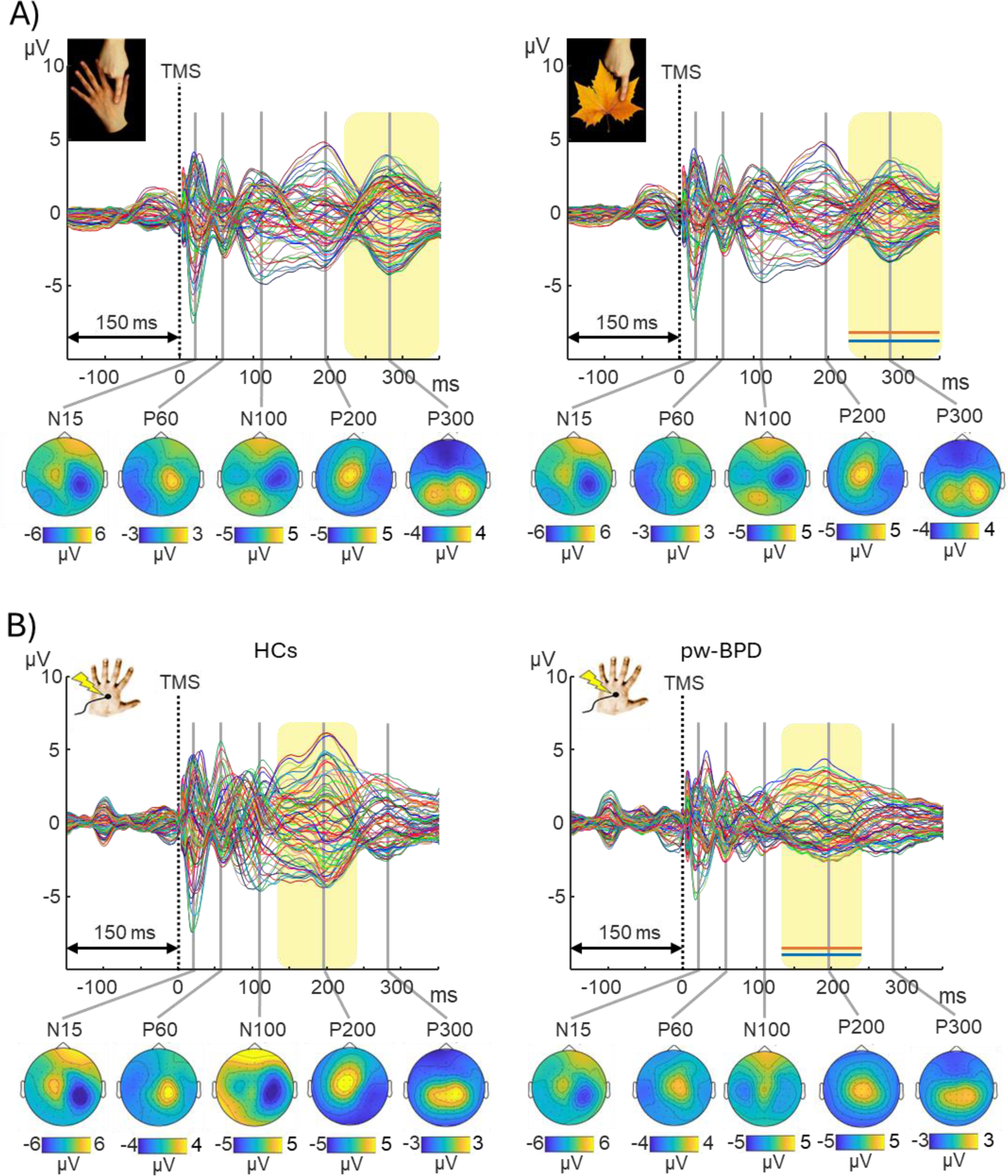
TEPs results in trials with ISI-150. **A)** Main effect of Stimulus: butterfly-plot TEPs generated 150 ms after visual touches on a hand (left panel) and on a leaf (right panel). **B)** Comparison between HCs (left panel) and pw-BPD (right panel) in TEPs generated 150 ms after real touches. Topographies (amplitude range in colorbar) are shown for main TEP components. Horizontal lines indicate the latencies of significant clusters (orange: positive; blue: negative).

Regarding TEPs recorded after real touches, exploratory analyses indicated differences between pw-BPD and HCs when TMS was delivered 150 ms after the real touch: while no significant clusters emerged in the preregistered analysis on ISI-20 trials (p>0.57), exploratory analyses on ISI-150 trials revealed two significant clusters (one positive, between 130 and 240 ms over left fronto-central electrodes, p=0.032; one negative, between 100 and 240 ms over right posterior central electrodes, p=0.034), revealing reduced TEPs amplitude in pw-BPD compared to HCs.

### Exploratory: ΔTEPs-ERPs peaks

Analyses on ΔTEPs-ERPs indicated a different connectivity pattern between HCs and pw-BPD, which was independent from the ISI or the visual stimulus (**Figure 5**; see **Figure S1** for an example of the subtraction process in visual touch trials in HCs). Peak amplitude and latency were extracted from two TEPs components in the ΔTEPs-ERPs, namely N15 and P60. A significant main effect of Group emerged for P60 (F(1,37)=6.92, p=0.012), with pw-BPD showing lower amplitude (mean ± SE: 3.19 ± 0.88 µV) compared to HCs (mean ± SE: 6.43 ± 0.86 µV). No other significant main effects or interactions were observed for the other peaks, either for the amplitude or the latency (summary of results in **Table S1**).

**Figure 5.**
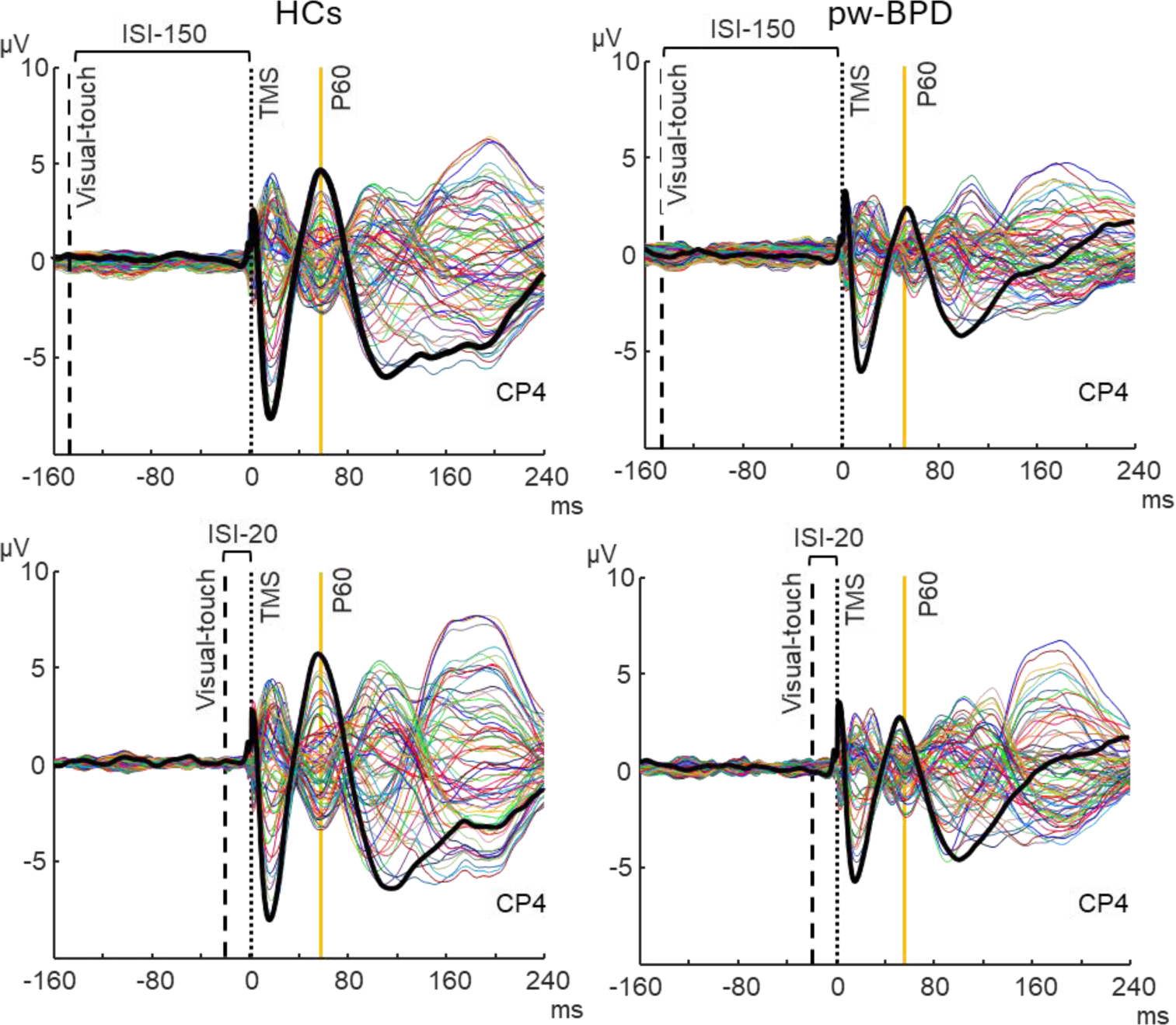
ΔTEPs-ERPs results. Butterfly plot of ΔTEPs-ERPs generated by S1-TMS delivered 150 ms (i.e., ISI-150, upper row) and 20 ms (i.e., ISI-20, lower row) after visual-touch trials on the hand in HCs (left panel) and pw-BPD (right panel). Black thick traces indicate channel CP4, which was selected for peak detection. Vertical orange lines indicate P60, for which we observed a main effect of Group in the Stimulus X ISI X Group rm-ANOVA.

## Discussion

In the present preregistered study pw-BPD and HCs went through a thorough investigation: they were evaluated and compared in the empathic abilities, in the behavioral performance in a task involving TaMS activity, and the neurophysiological measures obtained from TMS-EEG recording. The main findings show that pw-BPD reported significantly lower cognitive empathy, performed worse in terms of accuracy at the VTSC task involving TaMS activity, and displayed a different connectivity pattern in the TMS-EEG data, although it appeared to be not specific for TaMS but rather index of an alteration in the somatosensory network.

From a clinical point of view, BPD patients showed a moderate level of BPD symptoms, impulsiveness and depression. More than 80% of the sample reported self-harm behaviors.

Regarding empathic abilities, our findings suggest that pw-BPD have difficulties in understanding and imaging others’ perspective rather than feeling others’ sensations and experiencing their emotions. Specifically, the QCAE results show significantly lower levels in the cognitive domain in pw-BPD compared to HCs, but no difference in the affective domain, consistent with existent findings (Grzegorzewski et al., 2019). Although the results for the IRI on cognitive empathy did not reach statistical significance as in previous reports (Harari et al., 2010; Martin et al., 2017), they show a trend toward the same direction. Furthermore, in addition to previous studies, we provided normative values for the IRI scores (Maddaluno et al., 2022), showing that they were below or at the lower boundary for the normal range in half of the pw-BPD, which indicates that empathic impairment is a relevant aspect for BPD.

We hypothesized that such impairments in cognitive empathy in BPD could reflect alterations in the activity of mirror-like mechanisms in the somatosensory domain (Keysers et al., 2010). In the VTSC task used in the present study, seeing a touch on human body parts is expected to activate the TaMS, and the spatial incongruency between the seen touch and the felt one should interfere with the ability of reporting the side of the real touch (Banissy & Ward, 2007; Bolognini et al., 2013, 2014). Here, we report evidence of a greater interference effect in pw-BPD compared to HCs, indexed by a poorer performance at the VTSC task in terms of accuracy, only when seeing touch on body parts but not on objects, suggesting that the effect was specific for TaMS activity. This finding is intriguing, because existing behavioral evidence for mirror system alteration is scarce, with previous research showing more general impairments in inferring others’ mental states in pw-BPD (Dziobek et al., 2011; Harari et al., 2010; but see Mier et al., 2013; Preißler, Dziobek, Ritter, Heekeren, & Roepke, 2010). It is important to note that the direction of this result is different from what we expected, namely a reduced interference effect in pw-BPD. One possibility is that seeing touches on human body parts is particularly salient for BPD. Alternatively, their processing could be more demanding for BPD. Although these interpretations can only be speculative at this stage, it seems that the relationship between empathic abilities and TaMS activation is not linear as hypothesized, but rather more complex.

Differently, results on RTs in the VTSC task and on the subjective sensations did not show evidence of TaMS impairment under the conditions of the present study, but provided several quality checks on the VTSC task. Indeed, performance of both groups was influenced by the spatial congruency of the touching hand in relation to the real touch, indicating that the location of the visual stimulus was relevant to perform the task. Moreover, responses in visual-touch trials were faster than in unimodal tactile-only trials, showing a typical crossmodal facilitation (Macaluso & Maravita, 2010). In the HCs group, we did not observe either a significant difference in RTs between human- and object-directed visual-touches, or a significant relationship between cognitive empathy and VTSC performance. Such effects were previously reported on healthy participants (Bolognini et al., 2013, 2014), but, importantly, they were observed only after an experimental modulation of S1 activity by means of TMS or transcranial electrical stimulation, a crucial difference with the present paradigm. Therefore, our results on HCs should not be interpreted as in contrast with previous studies, and it may be hypothesized that the VTSC task is not sensitive enough to detect S1 mirror activity in baseline conditions. Considering that performance showed ceiling effects for HCs, future studies may test a more difficult version of the paradigm. Finally, the two-item questionnaire on the sensations induced by the VTSC task revealed a difference between hands and leaves block, indicating that participants (both HCs and pw-BPD) had a more intense feeling of being touched on their own hands during the hands block compared to the leaves block.

With respect to the neurophysiological underpinnings of TaMS activity, results obtained during the TMS-EEG recording did not show evidence of alterations in TaMS connectivity, according to the preregistered analyses. Nevertheless, they provided information both on the processing of the different visual-touch stimuli and their connectivity patterns, from ERPs and TEPs, respectively. To the best of our knowledge, the present study is the first one providing TEPs data in pw-BPD. Employing TMS during a task is known to enable the investigation of network-specific activity which is dependent on its functional status (Barchiesi et al., 2022; Jacquet & Avenanti, 2015; Silvanto, Muggleton, Cowey, & Walsh, 2007), while the simultaneous recording of EEG provides information on the spread of neural activation to brain areas effectively connected to the stimulated one (Bortoletto, Veniero, Thut, & Miniussi, 2015; Massimini et al., 2005; Zazio et al., 2021). Crucially, combining the two techniques in TMS-EEG recordings when participants are performing a task allows the investigation of brain connectivity in task-specific networks (Bortoletto et al., 2021; Morishima et al., 2009; Zazio et al., 2022). In both pw-BPD and HCs and in all conditions we successfully recorded clear TEPs components from S1 stimulation, namely N15, P60, N100, P200, P300, with topographical patterns overall similar to the ones described previously in HCs (Pisoni et al., 2018). When considering HCs and pw-BPD together, a difference between the obseration of human- and object-directed visual touches occurred at late latencies of TEPs, namely from 200 ms from the TMS pulse on. This finding suggests that TaMS activity involves S1 connection with distant areas. Moreover, it was observed when S1-TMS was delivered 150 ms - but not 20 ms - after touch onset, indicating that TaMS activation involves S1 in the high-order phase of the processing of visual-touches. This result supports previous studies (Bolognini et al., 2014; Pisoni et al., 2018) in identifying 150 ms as the timing of TaMS activation during touch observation.

Nevertheless, the present results from TEPs provide no evidence for a specific alteration S1-connectivity within TaMS in BPD, but they rather point to an impairment in the somatosensory network. Indeed, we did not observe any difference between pw-BPD and HCs in TEPs during touch observation, either when they were directed towards a body part or an object. Considering that TEPs were not recorded at rest but during the observation of touches, we also analyzed the difference between TEPs and ERPs (i.e., ΔTEP-ERP) to disentangle the contribution of the processing of the stimuli, as indexed by ERPs, from the S1-connectivity pattern, as indexed by TEPs. This procedure allowed the comparison between different ISIs without confounding factors, and the analysis of components’ peaks enabled us to include all relevant control conditions (i.e., group, stimulus and ISI) in a single statistical model. Results on ΔTEP-ERP peaks revealed a lower amplitude of P60 in pw-BPD compared to HCs irrespective of the ISI and of the visual stimulus, thus suggesting a general alteration in S1 connectivity in BPD. P60 may reflect a secondary activation of right sensorimotor areas (i.e., following the primary activation due to the TMS pulse): indeed, the topographical pattern of P60 shows a positive activity over right centro-parietal electrodes, and previous studies stimulating M1 has localized it in parietal areas (Zazio et al., 2021) and associated it with the somatosensory reafference of the motor-evoked potentials (Petrichella, Johnson, & He, 2017). Therefore, it may be hypothesized that the same circuit in the sensory-motor network could be activated following S1 stimulation. Moreover, in the study by Pisoni et al. (2018), the same component (called P50) has been attributed to vicarious S1 reactivity during the observation of human-directed touch. While extremely intriguing, this hypothesis remains speculative, as we did not observe a difference at this latency between the observation of touches on a body part and on an object. One possibility that cannot be ruled out is that P60 alteration reflects non-specific effects of the pharmacological treatments, as it does not interact with the ISI or the visual stimulus. However, it is worth noting that the resting motor threshold did not differ between pw-BPD and HCs, and this TEP difference was specific for P60 and not other TEP components, indicating that the effect on P60 cannot be due to general differences in cortical excitability. Consistently, results from TEPs recorded during real touches also indicate a perturbation of the somatosensory network in BPD. Indeed, the difference in TEPs between pw-BPD and HCs during real touches was present when TMS was delivered 150 ms but not 20 ms after the real touch (i.e., the timing of S1 activation following the somatosensory afference; (Cohen et al., 1991) and affected late components (i.e., 100 ms from the TMS pulse on), suggesting that the alteration affects the higher order stages of touch processing and S1 connections with distant areas, respectively.

On the other hand, exploratory results on ERPs point to an alteration in BPD selectively affecting the processing of visual-touches on body parts, thus index of TaMS activity: during the observation of touches (i.e., touchERPs), there was a tendency to reduced ERPs components (i.e., touchERPs) for hand-directed touches compared to object-directed touches in pw-BPD, an effect not present in HCs. Interestingly, this effect emerges approximately 200 ms after touch onset, which may suggest that such time interval might be better to probe TaMS connectivity alterations in BPD, compared to the 150 ms tested in the present study. Moreover, this finding seems to be specific for the observation of touches, as ERPs derived from the appearance of the static hand or of the leaf without any touch (i.e., baselineERPs) did not reveal discernible alterations between groups. Instead, baselineERPs confirmed that both in pw-BPD and HCs the two stimuli were processed differently both in pw-BPD and HCs, with the hand being the only one eliciting the N170 component which is typical for body parts (Kovács et al., 2006). However, the result of a possible alteration of the processing of human-directed visual touches in pw-BPD should be taken with caution as it did not reach the threshold for significance in the stimulus by group interaction, and therefore needs to be confirmed in future studies. Existing literature presents a variety of experimental paradigms: most of the evidence in this context comes from functional magnetic resonance imaging studies during the observation of visual stimuli with an emotional content, showing an increased or decreased pattern of activation in brain areas belonging to the mirror neuron system (Mier et al., 2013; Sosic-Vasic et al., 2019). In contrast, results from emotionally neutral stimuli, as in the present study, seem more subtle, with pw-BPD showing a trend for a different pattern of mu-desynchronization compared to HCs only at specific time points of an action observation task (Martin et al., 2017). Finally, no difference in ERPs were observed after real touches, consistent with previous studies that indicate no generalized dysfunction of basic somatosensation in pw-BPD (Malejko et al., 2018; Pavony & Lenzenweger, 2014). Interestingly, a recent study in which pw-BPD underwent a comprehensive psychophysical evaluation of different dimensions of touch perception showed impairments in tactile sensitivity, defined as the ability to detect a tactile stimulus on the skin, in the absence of deficits in tactile acuity (i.e., the ability to discriminate between two tactile stimuli presented in close proximity to one another; (Cruciani et al., 2023). Taken together, these findings converge in indicating that the impairment in the somatosensory system in pw-BPD is not “all or nothing”, but presents specific alterations which need to be further explored in future studies.

### Limitations

The present study has a few limitations. First, power analysis was based on previous data on empathy questionnaires, but for the VTSC task and TMS-EEG data, this was the first study involving BPD patients; so it may be underpowered for the between-subject comparisons, especially for connectivity analyses which considered all channels and time points. Data and results reported here can now be used for future sample size estimates. Moreover, in all the visual touch trials, both hands and leaves are touched by a hand, which may trigger additional mirroring mechanisms in sensorimotor cortices driven by the action observation network recruitment with a time course similar to the TaMS one (Guidali, Picardi, Franca, Caronni, & Bolognini, 2023; Valchev et al., 2016), even in the object control trials. To boost the contrast between touches on hands and objects, in future studies an object (e.g. a stick) may be used to represent the visual touch.

## Conclusion and future directions

To conclude, the present study supports previous findings on empathic dysfunction in BPD and provides novel insights on the characterization of TaMS, showing an impairment at the behavioral level. The evidence on S1-connectivity alteration provided by TEPs seemed not specific for the visuo-tactile mirror system, but rather reflecting a perturbation at higher order stages of the somatosensory network. Further research is needed to clarify the direction of the observed effects and the relationship with the deficits in empathic abilities.

## Authors’ contribution

AZ: conceptualization, investigation, data curation, methodology, formal analysis, visualization, writing - original draft, project administration, funding acquisition; supervision; CML: investigation, formal analysis, writing - original draft; AS: methodology, formal analysis, software, writing - review and editing; GG: conceptualization, investigation, software, visualization, writing - review and editing; EM: investigation, writing - review and editing; DL: investigation, writing - review and editing; SM: resources, writing - review and editing; RR: resources, writing - review and editing; NB: conceptualization, writing - review and editing; MB: conceptualization, writing - original draft.

## Acknowledgments

We would like to thank Matteo de Matola for his help in the analysis pipeline of TMS-EEG data.

## Funding

This project is supported by the Italian Ministry of Health (Bando Ricerca ‘Finalizzata 2019 – grant number SG-2019-12370473’ awarded to AZ and ‘Ricerca Corrente’).

## Conflict of interest statement

The authors declare that the research was conducted in the absence of any commercial or financial relationships that could be construed as a potential conflict of interest.

## Notes

https://osf.io/euymx/?view_only=eae250ff55e64665a052090bd1b41f9b

### Competing Interest Statement

The authors have declared no competing interest.

