## Supplementary material for "Investigating visuo-tactile mirror properties in Borderline Personality Disorder: a TMS-EEG study": Figure S1; Table S1

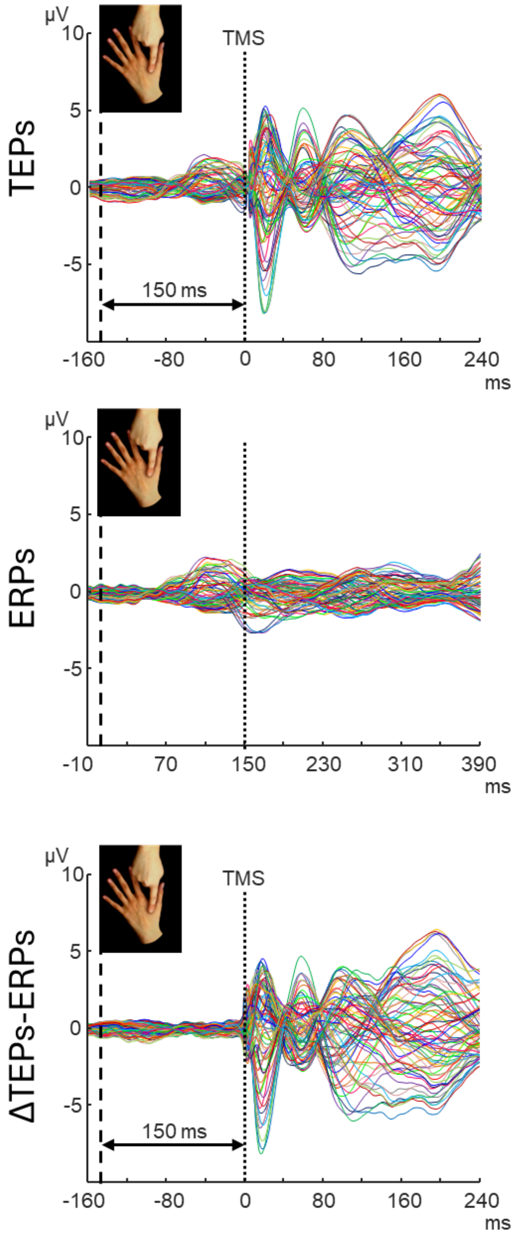


**Figure S1.** Butterfly plots showing ΔTEPs-ERPs subtraction process in hand-directed visuo-tactile trials in HCs. *Upper row:* TEPs recorded with ISI-150 (0 corresponds to TMS pulse). *Middle row:* ERPs from no-TMS trials (0 corresponds to visual touch onset; vertical dotted line at 150 ms shows the time point in which TMS would be delivered in case of TMS trials). *Lower row*: result of the ΔTEP-ERPs subtraction (0 corresponds to TMS pulse).

| | Amplitude | | | | | | Latency | | | | | |
|---|---|---|---|---|---|---|---|---|---|---|---|---|
| | N15 | | | P60 | | | N15 | | | P60 | | |
| | *F(1,37)* | *p* | $\eta^2 p$ | *F(1,37)* | *p* | $\eta^2 p$ | *F(1,37)* | *p* | $\eta^2 p$ | *F(1,37)* | *p* | $\eta^2 p$ |
| *Main effects* | | | | | | | | | | | | |
| Group | 1.520 | 0.225 | 0.039 | 6.920 | **0.012** | 0.158 | 0.102 | 0.751 | 0.003 | 2.370 | 0.132 | 0.060 |
| Stimulus | 1.360 | 0.251 | 0.025 | 0.000 | 0.991 | 0.000 | 0.493 | 0.487 | 0.013 | 0.822 | 0.370 | 0.022 |
| ISI | 0.006 | 0.936 | 0.000 | 0.425 | 0.518 | 0.011 | 0.017 | 0.898 | 0.000 | 0.019 | 0.891 | 0.001 |
| *Interactions* | | | | | | | | | | | | |
| Group X Stimulus | 0.940 | 0.338 | 0.025 | 0.138 | 0.713 | 0.004 | 1.157 | 0.289 | 0.030 | 1.589 | 0.215 | 0.041 |
| Group X ISI | 0.107 | 0.746 | 0.003 | 0.054 | 0.818 | 0.001 | 0.868 | 0.357 | 0.023 | 4.765 | 0.035 | 0.114 |
| Stimulus X ISI | 0.062 | 0.805 | 0.002 | 0.007 | 0.935 | 0.000 | 4.701 | 0.037 | 0.113 | 0.348 | 0.559 | 0.009 |
| Group X Stimulus X ISI | 0.412 | 0.525 | 0.011 | 0.238 | 0.628 | 0.006 | 3.738 | 0.061 | 0.092 | 0.219 | 0.642 | 0.006 |

**Table S1.** Results of the Stimulus X ISI X Group rm-ANOVA on ΔTEPs-ERPs peaks (N15 and P60 amplitude and latency). Significant p value (following multiple comparisons correction) for the main effect of Group on P60 amplitude is highlighted in bold.
